# Measuring the Stoichiometry of Microbial Parasite Infrapopulations One Cell at a Time using Energy Dispersive Spectroscopy

**DOI:** 10.1101/2024.01.11.575226

**Authors:** Charlotte F Narr, Scott Binger, Erin Sedlacek, Bianca Anderson, Grace Shoemaker, Adrienne Stanley, Madison Stokoski, Ed Hall

## Abstract

1. Progress in the field of ecological stoichiometry has demonstrated that the outcome of ecological interactions can often be predicted *a priori* based on the nutrient ratios (e.g., carbon: nitrogen: phosphorus, C:N:P) of interacting organisms. However, the challenges of accurately measuring the nutrient content of active parasites within hosts has limited our ability to rigorously apply ecological stoichiometry to host-parasite systems. Traditional nutrient analyses require high parasite biomasses, often preventing individual-level analyses. This prevents researchers from estimating variation in the nutrient content of individual parasites within a single host infrapopulation, a critical factor that could define how the ecology of the parasite affects the host-parasite interaction.
2. Here, we explain how energy dispersive technology, a technique currently used to measure the elemental content of free-living microbes, can be adapted for parasitic microbial infrapopulations. We demonstrate the power of accurately quantifying the biomass stoichiometry of individual microbial parasites sampled directly from individual hosts.
3. Using this approach we show that the stoichiometric composition of two microbial parasites capable of infecting the same host are stoichiometrically distinct and respond to host diet quality differently. We also demonstrate that characteristics of the stoichiometric trait distributions of these infrapopulations were important predictors of host fecundity, a proxy for virulence in this system, and better predictors of parasite load than the mean parasite stoichiometry or our parasite and diet treatments alone.
4. EDS provides a rigorous tool for applying ecological stoichiometry to host-parasite systems and enables researchers to explore the nutritional physiology of host-parasite interactions at a scale that is more relevant to the ecology and evolution of the system than traditional nutrient analyses. Here we demonstrate that this level of resolution provides useful insights into the diet-dependent physiology of microbial parasites and their hosts. We anticipate that this improved level of resolution has the potential to elucidate a range of eco-evo interactions in host-parasite systems that were previously unobservable.

## Introduction

Multiple frameworks use concepts from the field of ecological stoichiometry to mechanistically predict the outcome of host-parasite interactions by comparing the stoichiometry of parasite biomass to that of the host and host’s environment (Aalto, Decaestecker, & Pulkkinen, 2015; Bernot & Poulin, 2018; Sanders & Taylor, 2018; Vannatta & Minchella, 2018). While this logic is intuitive and engaging, methodological challenges associated with testing these frameworks currently limit their utility. For example, a recent metanalysis could not explain the direction or magnitude of the effects of host diet quality on parasite virulence (Pike, Lythgoe, & King, 2019). Directly quantifying the nutritional ecology of host-parasite interactions could transform our understanding of how parasites influence their hosts’ population dynamics (Ebert, Lipsitch, & Mangin, 2000), evolution (Lively, 1987), communities (Wood et al., 2007), and ecosystems (Kuris et al., 2008).

One issue limiting the application of a stoichiometric mass balance approach to host-parasite systems is that data on parasite biomass stoichiometry is sparse and typically relies on traditional nutrient analyses that require high quantities of parasite biomass. These biomass requirements make it much easier to measure the stoichiometry of macroparasites than microbial parasites. As a result, the handful of studies that have measured the stoichiometry of parasites of animals have measured those of macroparaistes (e.g. Bernot, 2013; Narr & Krist, 2015; Paseka & Grunberg, 2018). This focus on macroparasites limits our ability to test basic hypotheses about the nutritional physiology of parasites given macroparasites often have complex life cycles that make it difficult to manipulate, or even quantify, the nutrient content of the host or its diet. Conversely, the simple life cycle and rapid growth of microbial parasites make them excellent model systems for testing fundamental hypotheses related to disease (Ebert, 2005), their role in nutrient cycling (Narr & Frost, 2015), and host resource-disease interactions (Civitello, Allman, Morozumi, & Rohr, 2018). Despite these advantages, we are aware of only a single study measuring the stoichiometry of animal microbial parasites in *vivo*, and it required integrating the biomass from 50 hosts to achieve sufficient biomass for a single, quantifiable replicate (Frost, Ebert, & Smith, 2008).

The application of stoichiometric frameworks to host-parasite interactions also currently omits an important aspect of parasite ecology by representing the stoichiometry of a parasite within a host as a single mean value. This limits our understanding of the ecology and evolution of these systems because parasites typically exist as infrapopulations comprised of multiple individuals within hosts (Poulin, 2007). As individuals, these constituents of an infrapopulation likely vary in their phenotypes, physiology, and stoichiometric requirements, and this variation likely shapes competition for nutrients with the host. We propose that describing an infrapopulation’s stoichiometric variation by quantifying its distribution of stoichiometric traits (biomass C:N, C:P and N:P ratios) could help predict outcomes of the host-parasite interaction.

Here, we present a method for analyzing the biomass stoichiometry of individual microbial parasites (spores) in individual hosts using energy dispersive spectroscopy (EDS). This workflow was recently re-established to characterize the stoichiometric trait distributions of freshwater planktonic communities (Manzella, Geiss, & Hall, 2019). Here, we explain how this technique can be adapted to quantify the stoichiometric trait distributions of parasite infrapopulations *in vivo*. We then present metrics that describe the phenotypic trait space (e.g. richness, evenness, divergence, etc.) occupied by constituents of an infrapopulation. We apply this approach to a well-studied *Daphnia*-microbial parasite system and compare the effects of host diet quality on the biomass stoichiometry of two microbial parasites capable of infecting the same host. We anticipate that this approach has the potential to elucidate fundamental ecological mechanisms of a wide range of host-parasite dynamics for diverse host-parasite systems.

## Methods

### Preparation of individual Daphnia samples for EDS analysis

We measured the stoichiometric ratios of microbial infrapopulations in *D. magna*. First, we created homogenized solutions of individual *Daphnia* by rinsing and then crushing them in ultrapure water. We adjusted the volume of water in our samples to attain ∼200-500 spores per grid and minimize spore overlap with other material from *Daphnia* bodies. Then, we pipetted 2 μl of each solution onto a nickel transmission electron microscope (TEM) grid with a silicon monoxide film (SF200-Cu, Electron Microscopy Supplies) and allowed it to dry at room temperature.

### EDS Analyses of spores

We visually identified individual spores based on morphology and then traced them with a within system drawing tool on a scanning electron microscope (JEOL JSM-6500F) equipped with an 80 mm^2^ Oxford X-max EDS detector following the methods of Manzella et al. (2019) (Figure 1). We used a grid holder that functions as a STEM converter (JU2002890, JEOL) to separate the TEM grid from the metal holder and avoid complications that have limited the use of SEM for biological samples in the past (Segura-Noguera, Blasco, & Fortuño, 2012). The entirety of the identified area was then rastered with the ion beam for ∼40 seconds with a probe current of 11 and a beam aperture of 3. These settings allowed for excitation of the cellular biomass and maximized signal to noise ration without completely destroying each spore. To create spectra characteristic of the elemental composition of each spore, X-rays were then detected by the EDS arm of the instrument. We also created control spectra by exciting a cell free area of analogous size that contained only the SiO_2_ film and any associated detritus (Figure 1).

**Figure 1:**
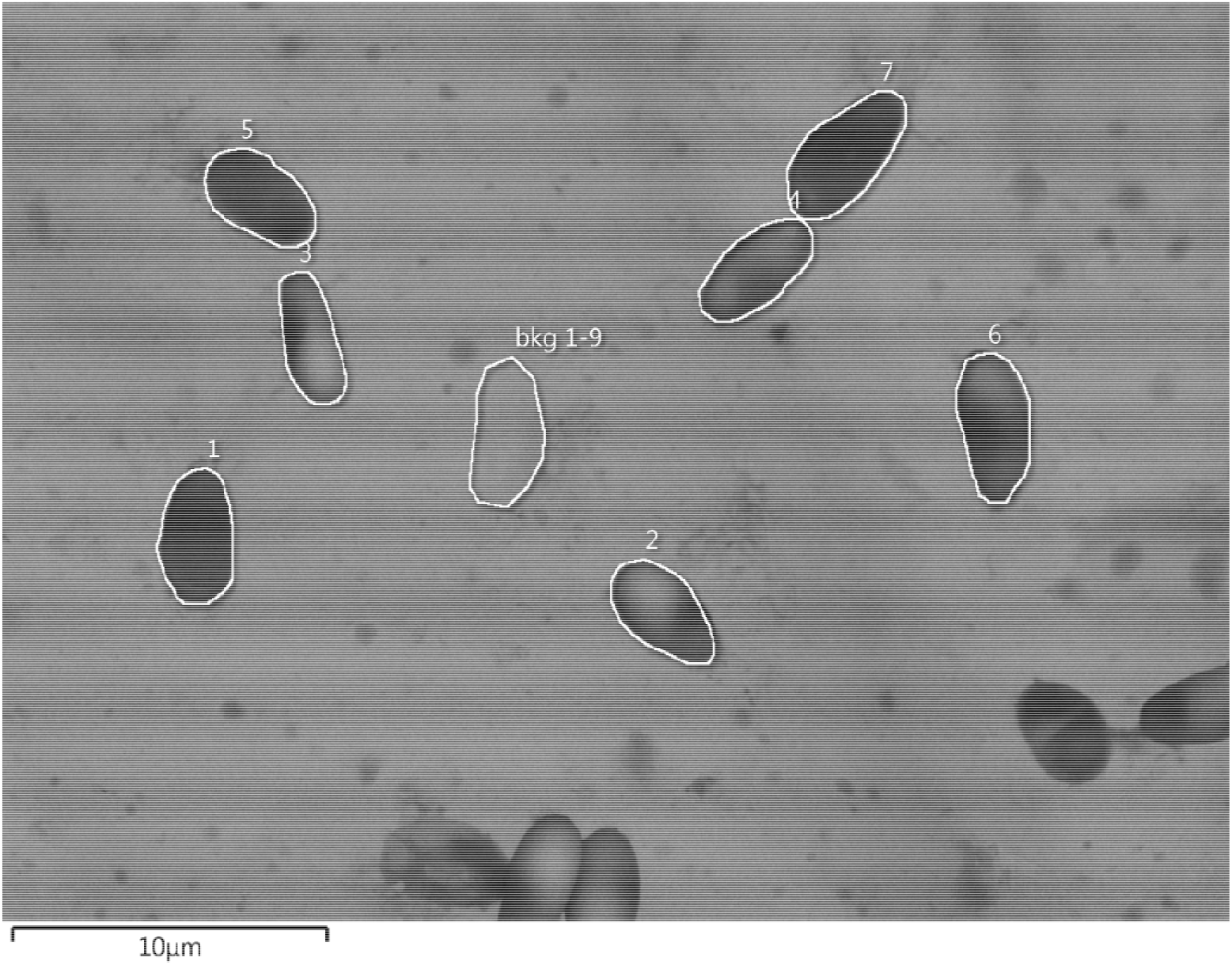
Scanning electron micrograph image of *Hamiltosporidium tvarminnensis* spores from *Daphnia* raised on a high P diet. The white lines highlight spores that were selected for EDS analyses and a single background area (bkg1-9) that was representative in size of cells 1-9 (cells 8 and 9 are not pictured). The entire area that is highlighted here was excited for analysis.

### EDS Data processing and analysis

We analyzed each spectrum with the freely-available NIST-based DTSA II (Desktop Spectral Analyzer) software program Jupiter. This program allowed us to strip the background (continuum) radiation, subtract the contribution of the silica film from each spectra, and quantify the total counts within pre-selected ranges derived from biochemical standards (ATP, ADP and Acetyl-CoA, Manzella,Geiss and Hall 2019). We removed the control spectra from the spectra containing microbial parasite, and these values were multiplied by the previously established calibration constants (Manzella, Geiss and Hall 2019) that convert EDS output (counts per second, cps) to the amount of C, N, and P (fm) within each cell. Here we report molar ratios of C, N and P derived from these atomic cell quotas.

### Experimental Procedure

Additional details regarding experimental procedure are provided in Appendix A. Briefly, we infected *Daphnia magna* with the microsporidian parasite *Hamiltosporidium tvaerminnensis* and the bacterial parasite *Pasteuria ramosa. Daphnia* neonates (<24 h old) were taken from the 4-5^th^ broods of mothers maintained under high food quality and quantity conditions, raised in COMBO (Kilham et al 1998) and fed a controlled quantity of one of two diet treatments (*Scendesmus obliquus* with C:P ratios of approximately 100 and 600, Appendix 1, Table 1) every other day. Neonates were collected every other day, stored, and then counted to assess individual fecundity.

*Daphnia* were raised until day 28, measured, and then prepared for EDS analysis. We estimated individual spore loads from the homogenized *Daphnia* solution described above, and then we selected two *Daphnia* infected with *H. tvaerminensis* and three infected with *P. ramosa* for EDS to represent the range of spore loads within each treatment. Spore loads were estimated by dividing the average total number of spores in each sample (estimated from 3 - 16 μl aliquots on a hemocytometer) by the *Daphnia*’s estimated weight (calculated according to Ku et al., 2022).

### Statistical Analysis

To determine if parasite type or diet influenced spore stoichiometry, we used linear mixed models that included parasite type, diet and their interaction as fixed effects and individual host as a random effect. Ratios were log-transformed to avoid biases in interpretation (Isles, 2020). Models revealing significant interactions between parasite type and diet were followed by separate linear mixed models for each parasite with diet as a fixed effect and individual host as a random effect. Satterthwaite’s method was used to assess significance.

We calculated the functional richness, evenness and divergence of each infrapopulation based on spore C:N, C:P, and N:P ratios using the TPD package in R (Carmona et al. 2019). Then, we compared our ability to explain the spore load and fecundity of each *Daphnia* with univariate models comprised of: each distribution metric, the mean biomass stoichiometry of the infrapopulations, diet, parasite treatment, or a null model. We used Akaike’s information criteria corrected for small sample sizes (AICc). Models with a delta AICc < 2 were considered equivalent. We modeled spore load with negative binomial models and fecundity (log-transformed to achieve normally distributed errors) with linear models. All statistics were run in R version 4.3.0.

## Results

For both diets, spores from *P. ramosa* infrapopulations had higher C:P and N:P ratios than spores from *H. tvaerminnensis* infrapopulations (C:P: t = 5.621, p<0.001, N:P: t = 5.456, p < 0.001, Figure 2). However, an interaction between diet and infection (t = -2.48, p = 0.032) suggested that diet influenced *P. ramosa* and *H. tvaerminnensis* infrapopulation C:P ratios in different ways. Specifically, low P diets decreased the C:P ratios (t = -2.598, p = 0.040) of *P. ramosa* spores but increased the C:P ratios of *H. tvaerminnensis* spores (t = 2.053, p = 0.041) relative to high P diets.

**Figure 2:**
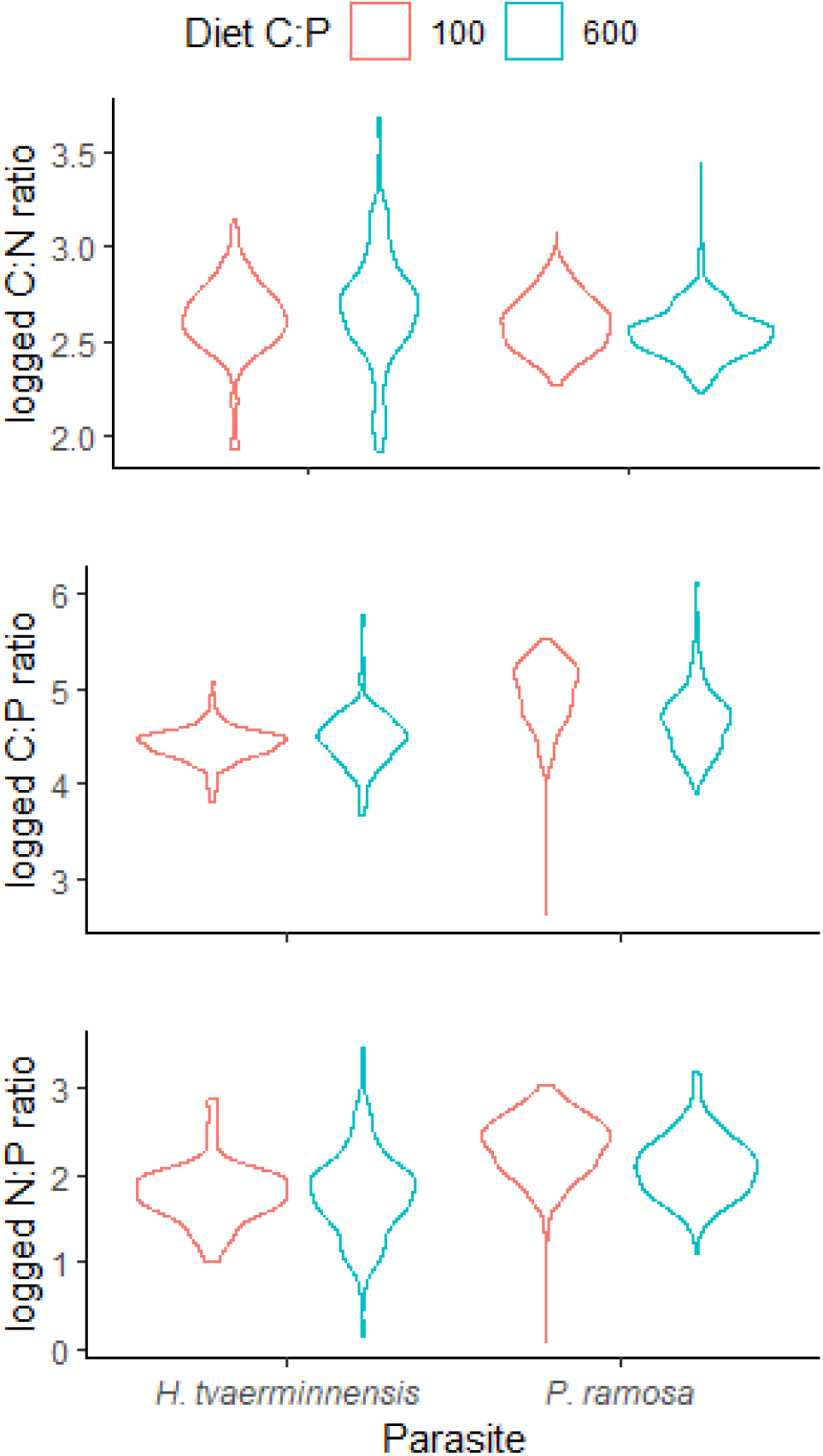
Log transformed stoichiometric ratios of spores from infrapopulations of *Hamiltosporidium tvaerminnensis* and *Pasteuria ramosa* in daphnid hosts raised on diets with C:P ratios of 100 (red) and 600 (blue).

Model selection indicated that C:N divergence was the best predictor of spore load (Appendix, Table 2). Both C:P divergence and parasite treatment were the best performing models for predicting fecundity (Appendix 1, Table 3). Specifically, *Daphnia* infected with *H. tvaerminnensis* had higher fecundity than those infected with *P. ramosa* (t=3.678_(1,8)_, p = 0.0062). However, spore load and fecundity also declined with infrapopulation divergence in C:N and C:P, respectively (Spore load: Z = - 3.82, p < 0.001, Fecundity: F = 10.91_(1,8)_, p = 0.011, Figure 3).

**Figure 3:**
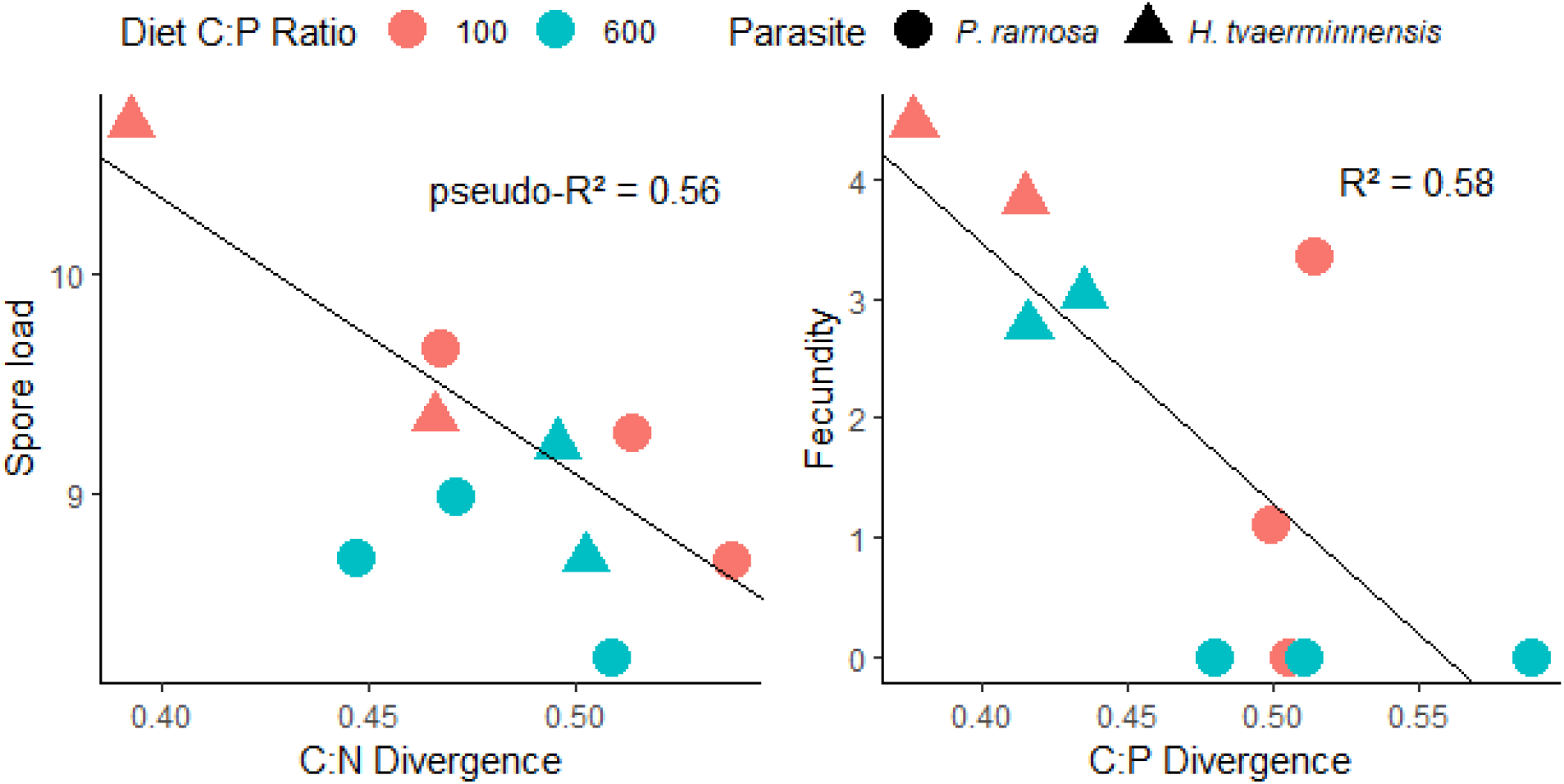
Log transformed spore load (million spores/μg dry weight) and fecundity (total offspring produced over 28 days) of *Daphnia* raised on diets with C:P ratios of 100 (red) and 600 (blue) infected with infrapopulations of *P. ramosa* (circles) or *H. tvaerminnensis* (triangles).

## Discussion

Here we show how EDS can be used to rigorously quantify both the mean nutrient ratios and stoichiometric trait distribution of microbial parasite infrapopulations within individual hosts. A major advantage of this approach is that it takes into account the ecology of microbial parasites and conceptually and methodologically treats the infrapopulation as a group of individuals rather than a single entity. This provides the opportunity to query the ecology of microbial parasites and identify fundamental ecological mechanisms that underlie previously undescribed host-parasite dynamics. Below, we highlight considerations for adapting this method to other host-parasite systems and discuss the implications of our findings in the *Daphnia*-microbial parasite system. The lack of comparable data in the literature highlights how difficult it is to test fundamental hypotheses applying ecological stoichiometry to host-parasite systems using traditional nutrient analyses.

### Applying EDS to other host-parasite systems

There are a few considerations for effective use of EDS to measure the stoichiometry of microbial parasites. First, EDS is likely to be most appropriate for microbial parasites that are close to 1 μm in size. Generating TEM grids with appropriate parasite densities (we suggest ∼200-500/ grid) will enable researchers to reduce search time and minimize clumping of cells. We also suggest some trial and error to minimize overlap between parasites and host tissues on grids. However, in systems where parasites achieve smaller loads within a single host, it may be necessary to sample a specific tissue or infection site before homogenizing the sample. In its current form, our method is likely to be most appropriate for microbes in cell culture or in hosts with simple immune systems. In other systems (e.g. intracellular pathogens of vertebrates), separating *in vivo* parasites from host tissues will likely require additional method development or alternatively, preferential use of cell culture infection models. Finally, an ability to identify individual parasites from SEM images is necessary for this method. The microbial parasite that we studied have characteristic morphologies that facilitated our ability to easily identify them and distinguish them from similarly sized particles within SEM images (Figure 1).

### Insights from EDS on parasitic infrapopulations of Daphnia

Determining if and how parasite identity influences its stoichiometry is a critical first step in applying ecological stoichiometry to host-parasite systems, but stoichiometric comparisons across parasites reared in controlled environments are, to our knowledge, non-existent. The application of this method to the *Daphnia*-microbial parasite system shows how these comparisons can provide a novel and mechanistic understanding of host-parasite systems. Our method revealed that individuals from *P. ramosa* infrapopulations had higher C:P and N:P ratios than those from *H. tvaerminnensis* infrapopulations, regardless of host diet (Figure 2). These trends were consistent with previous work showing that *Daphnia* infected by *P. ramosa* had higher C:P ratios and recycled phosphorus more quickly than *Daphnia* infected by *H. tvaerminnensis* (Narr & Frost, 2016). By measuring the stoichiometry of the parasite itself, our approach provided evidence that these shifts in host nutrient recycling may have been due to the accumulation of nutrients in parasite biomass. Some evidence supporting the hypotheses that parasites differ in their nutrient content (Paseka & Grunberg, 2019) and that infection alters nutrient recycling form hosts has been obtained from other host-parasite systems (Bernot, 2013; Chodkowski & Bernot, 2017; Mischler, Johnson, McKenzie, & Townsend, 2016; Narr & Krist, 2015; Paseka & Grunberg, 2019), but this work was limited to helminths (macroparasites) from wild hosts.

While data from wild hosts is valuable, our results suggest that observing the stoichiometry of parasites under controlled conditions is also important, in part, because host diet can influence parasite stoichiometry. In our application, low P host diets increased the C:P ratios of the microsporidian infrapopulation and, counterintuitively, decreased the C:P ratios of the bacterial infrapopulation. The unexpected response of the bacterial parasite to host nutrient limitation highlights the need for additional work examining the causes and consequences of host diet manipulations on the stoichiometric ratios of multiple parasite types. To date, we are aware of only one study that has quantified the effect of host diet on parasite stoichiometry in animals (Narr and Krist, 2015), and one that evaluated a similar relationship in phytoplankton hosts (manipulating nutrient availability in the environment of phytoplankton hosts, Frenken et al., 2017). However, the constraints of traditional nutrient analyses limited the resolution of previous work to the single mean stoichiometric values for individual or pooled hosts. As indicated by our model comparison, this limits our ability to investigate the source of this variation and quantify potential tradeoffs associated with this variation across parasites and environments.

Our method allowed us to measure stoichiometric variation within infrapopulations, and our analyses suggest that characterizing this variation can provide more information about variation in spore loads and virulence than the mean of the distribution. This suggests that traditional nutrient analyses (which, at best, quantify the mean nutrient content of a parasite within a host and could include contamination from host biomass) limit the ability to infer the role of the parasite’s nutrient content in shaping parasite or host success. In our data, divergence in C:N ratios within infrapopulations was the top predictor of spore load, and infrapopulations with greater C:N ratio divergence had smaller spore loads. Similarly, divergence in C:P ratios was a top predictor of fecundity, and infrapopulations with greater C:P ratio divergence had lower fecundities. These relationships suggest that divergence of stoichiometric traits may be costly to both the parasite and host. If so, these trends should become more evident over longer timescales. Investigating this possibility and many other fundamental questions about the role of parasite nutrient content in shaping the ecology and evolution of parasitism should be greatly facilitated by the ease with which this method can be applied to host-parasite systems that are easy to manipulate in the lab.

## Notes

### Competing Interest Statement

The authors have declared no competing interest.

