## Appendix A for "Measuring the Stoichiometry of Microbial Parasite Infrapopulations One Cell at a Time using Energy Dispersive Spectroscopy"

Appendix 1

*Additional Details on Experimental Procedure*

We infected a single *Daphnia magna* clone (OER 3-3) with the microsporidian parasite *Hamiltosporidium tvaerminnensis*, which resides in the host’s gut and ovaries (Vizoso & Ebert, 2004)and the bacterial parasite *Pasteuria ramosa*, which infects the hemolymph of *Daphnia* (Ebert et al., 1996). Daphnia neonates (<24 h old) were taken from the 4-5^th^ broods of mothers maintained under high food quality and quantity conditions, raised individually in 20 mls (days 0-6) and then 40 mls (days 6-28) of COMBO (Kilham et al 1998) and fed 2 (days 0 and 2), 4 (day 4) and 8 (days 6-28) mg C/L every other day. Starting on day six, we transferred *Daphnia* to new media every 4 days. All neonates were collected and stored in 90% ethanol every other day, and then counted to assess individual fecundity. We created 2 diets of *Scendesmus obliquus* by spiking the algal media of continuously grown cultures with NaH_2_PO_4_, and then mixing these cultures to create diets with C:P ratios of approximately 100 and 600 (appendix 1, table1).

We obtained *Daphnia* that were vertically infected with *H. tvaerminnensis* by collecting the 4-5^th^ brood of *H. tvaerimnensis* infected *Daphnia*. One generation prior to the experiment, these brood mothers were taken as neonates from stock uninfected brood moms and horizontally infected by exposure to the crushed bodies of *H. tvaerminnensis*-infected *Daphnia*. To create *P. ramosa* infections, uninfected neonates were exposed to a dose of 100,000 spores via a solution of homogenized infected *Daphnia* bodies from days 0-6. *H. tvaerminnensis*-infected *Daphnia* were exposed to the same dose of homogenized uninfected *Daphnia*. *Daphnia* were raised until day 28, measured, and then prepared for EDS analysis. We estimated individual spore loads from the homogenized Daphnia solution described above, and then selected *Daphnia* for EDS to represent the range of spore loads within each treatment. Spore loads were estimated by dividing the average total number of spores in each sample (estimated from 3 - 16 µl aliquots on a hemocytometer) by the *Daphnia*’s estimated weight (Ku, et al., 2022). We selected two *Daphnia* infected with *H. tvaerminensis* and three infected with *P. ramosa* from each diet for EDS analysis.

Table 1: Mean_(SE)_ for C:N, C:P, N:P and %C, P, and N of algal diets (identified by their desired C:P ratio). Values represent the post-mixing composition of diets fed to animals throughout the experiment.

| **Diet** | **C:N** | **C:P** | **N:P** | **%C** | **%N** | **%P** |
| --- | --- | --- | --- | --- | --- | --- |
| **100** | 10.2_(0.4)_ | 83.4_(7.6)_ | 8.3_(0.8)_ | 48.7_(0.6)_ | 5.6_(0.2)_ | 1.6_(0.2)_ |
| **500** | 12.3_(0.4)_ | 516.0_(68.2)_ | 42.0_(5.6)_ | 53.3_(0.9)_ | 5.1_(0.1)_ | 0.3_(0.04)_ |

Table 2: Model selection for univariate negative binomial models predicting the spore load of individual hosts based on characteristics of the stoichiometric trait distributions of their parasitic infrapopulations or on diet or infection treatments. Number of parameters (K), change in AIC_C_ compared to the best-ranked model (ΔAIC_C_), Akaike model weights (*W*), and log likelihood estimate (*LL*) for top models (ΔAIC_C_ < 2) are shown in bold.

| Model Predictor | K | ΔAICc | *W* | *LL* |
| --- | --- | --- | --- | --- |
| **C:N divergence** | **3** | **0.00** | **0.77** | **-97.03** |
| Diet Quality | 3 | 5.48 | 0.05 | -99.77 |
| N:P evenness | 3 | 5.81 | 0.04 | -99.93 |
| N:P divergence | 3 | 6.49 | 0.03 | -100.27 |
| C:P divergence | 3 | 6.86 | 0.03 | -100.46 |
| Null | 2 | 7.10 | 0.02 | -102.72 |
| Infection | 3 | 7.90 | 0.01 | -100.98 |
| C:P evenness | 3 | 8.19 | 0.01 | -101.12 |
| C:N evenness | 3 | 8.20 | 0.01 | -101.13 |
| C:P richness | 3 | 9.10 | 0.01 | -101.58 |
| N:P richness | 3 | 10.64 | 0.00 | -102.35 |
| N:P mean | 3 | 10.91 | 0.00 | -102.48 |
| C:N richness | 3 | 10.97 | 0.00 | -102.51 |
| C:P mean | 3 | 10.99 | 0.00 | -102.52 |
| C:N mean | 3 | 11.22 | 0.00 | -102.64 |

Table 3: Model selection for univariate linear models predicting the log transformed fecundity of individual hosts based on characteristics of the stoichiometric trait distributions of their parasitic infrapopulations or on diet or infection treatments. Number of parameters (K), change in AIC_C_ compared to the best-ranked model (ΔAIC_C_), Akaike model weights (*W*), and log likelihood estimate (*LL*) for top models (ΔAIC_C_ < 2) are shown in bold.

| Model Predictor | K | ΔAICc | *W* | *LL* |
| --- | --- | --- | --- | --- |
| **Infection** | **3** | **0.00** | **0.41** | **-14.67** |
| **C:P divergence** | **3** | **1.30** | **0.22** | **-15.32** |
| C:N mean | 3 | 2.15 | 0.14 | -15.75 |
| C:N richness | 3 | 4.47 | 0.04 | -16.91 |
| C:P richness | 3 | 4.63 | 0.04 | -16.99 |
| Null | 2 | 5.61 | 0.02 | -19.62 |
| N:P divergence | 3 | 5.81 | 0.02 | -17.58 |
| C:N evenness | 3 | 5.86 | 0.02 | -17.61 |
| C:P evenness | 3 | 6.36 | 0.02 | -17.85 |
| N:P evenness | 3 | 6.68 | 0.01 | -18.01 |
| N:P mean | 3 | 7.00 | 0.01 | -18.17 |
| C:N divergence | 3 | 7.35 | 0.01 | -18.35 |
| N:P richness | 3 | 7.52 | 0.01 | -18.43 |
| Diet Quality | 3 | 8.12 | 0.01 | -18.73 |
| C:P mean | 3 | 8.42 | 0.01 | -18.88 |
